# Dynamics and quantitative contribution of the aminoglycoside 6′-*N*-acetyltransferase type Ib [AAC(6′)-Ib] to amikacin resistance

**DOI:** 10.1101/2023.09.05.556435

**Authors:** Ophélie d’Udekem d’Acoz, Fong Hue, Tianyi Ye, Louise Wang, Maxime Leroux, Lucila Rajngewerc, Tung Tran, Kimberly Phan, Maria S. Ramirez, Walter Reisner, Marcelo E. Tolmasky, Rodrigo Reyes-Lamothe

## Abstract

Aminoglycosides are essential components in the available armamentarium to treat bacterial infections. The surge and rapid dissemination of resistance genes strongly reduce their efficiency, compromising public health. Among the multitude of modifying enzymes that confer resistance to aminoglycosides, the aminoglycoside acetyltransferase AAC(6′)-Ib is the most prevalent and relevant in the clinical setting as it can inactivate numerous aminoglycosides, such as amikacin. Although the mechanism of action, structure, and biochemical properties of the AAC(6′)-Ib protein have been extensively studied, the contribution of the intracellular milieu to its activity remains unclear. In this work, we used a fluorescent-based system to quantify the number of AAC(6′)-Ib per cell in *Escherichia coli,* and we modulated this copy number with the CRISPR interference method. These tools were then used to correlate enzyme concentrations with amikacin resistance levels. Our results show that resistance to amikacin increases linearly with a higher concentration of AAC(6′)-Ib until it reaches a plateau at a specific protein concentration. *In vivo* imaging of this protein shows that it diffuses freely within the cytoplasm of the cell, but it tends to form inclusion bodies at higher concentrations in rich culture media. Addition of a chelating agent completely dissolves these aggregates and partially prevents the plateau in the resistance level, suggesting that AAC(6′)-Ib aggregation lowers resistance to amikacin. These results provide the first step in understanding the cellular impact of each AAC(6′)-Ib molecule on aminoglycoside resistance. They also highlight the importance of studying its dynamic behavior within the cell.

**Importance:** Antibiotic resistance is a growing threat to human health. Understanding antibiotic resistance mechanisms can serve as foundation for developing innovative treatment strategies to counter this threat. While numerous studies clarified the genetics and dissemination of resistance genes and explored biochemical and structural features of resistance enzymes, their molecular dynamics and individual contribution to resistance within the cellular context remain unknown. Here, we examined this relationship modulating expression levels of AAC(6′)-Ib, an enzyme of clinical relevance. We show a linear correlation between copy number of the enzyme per cell and amikacin resistance levels up to a threshold where resistance plateaus. We propose that at concentrations below the threshold, the enzyme diffuses freely in the cytoplasm but aggregates at the cell poles at concentrations over the threshold. This research opens promising avenues for studying enzyme solubility’s impact on resistance, creating opportunities for future approaches to counter resistance.

## Introduction

The emergence and rapid dissemination of antibiotic resistance genes are among the biggest threats to global health (1, 2). Infections caused by antibiotic-resistant bacteria are rising in hospital and community settings. This growing trend is undermining treatment options and posing a significant risk to the success of medical and dental procedures that rely on preventing bacterial contamination (1). The current number of fatal infections due to resistance is estimated at hundreds of thousands per year, and the number could grow in the future (2). New drugs are urgently needed, but despite calls for action by diverse world organizations, the number of potential new drug candidates remains critically low (1–3). Two major therapeutic strategies have been proposed to deal with this crisis: (1) designing new antibiotics or (2) inhibitors of resistance that can be combined with existing antimicrobials (4). The success of these novel therapies depends on a detailed understanding of the different cellular and molecular mechanisms by which bacteria resist antibiotics (4).

Aminoglycosides are broad-spectrum antibiotics that interfere with normal protein synthesis by binding to the 16S rRNA. While not all classes of aminoglycosides bind to identical sites of the 16S rRNA, in all cases the A site (the ribosome’s decoding center) undergoes a conformational change to one resembling the closed state, formed after interacting with the cognate tRNA and mRNA (5). Consequently, the proofreading capabilities of the ribosome are reduced or eliminated, resulting in high levels of mistranslation (6, 7). However, despite the remarkable advances in understanding the mechanism of action of aminoglycosides, there is still much to learn about the nature of the translation errors generated by these drugs (8). Aminoglycosides have become less popular after being broadly utilized for decades due to increased resistance and relatively high toxicity. However, the requirement for effective medications and the need to develop less harmful semisynthetic variations to overcome the most prevalent resistance mechanisms have revitalized interest in their use (9, 10).

Amikacin, one of the most successful semisynthetic aminoglycosides (11), is refractory to most aminoglycoside modifying enzymes, which through acetylation, phosphorylation, or nucleotidylation, are the most common causes of resistance in clinical settings. Unfortunately, amikacin is a substrate for aminoglycoside 6′-*N*-acetyltransferase type I enzymes [AAC(6′)-I], which promote inactivation through acetylation at the 6’ amine group of the antibiotic (7, 11). Among *aac(6′)-I* genes, *aac(6′)-Ib* is the most clinically relevant, being found in the vast majority of multiple aminoglycoside-resistant Gram-negative isolates (12). The *aac(6′)-Ib* gene is located within plasmids and chromosomes as part of integrons, transposons, genomic islands, and other genetic structures that facilitate its dissemination at the molecular and cellular level, reaching virtually all Gram-negative bacteria (13). Here, we used the *aac(6’)-Ib* gene located in Tn*1331*, a transposable element present in pJHCM1, a plasmid isolated from a clinical *K. pneumoniae* strain (14, 15). While considerable advances have been made in the understanding of the dissemination mechanisms of *aac(6′)-Ib* (7, 16, 17), structural characteristics of the enzyme (18, 19), and its specificity properties (13, 20–22), multiple other aspects remain to be elucidated.

A significant gap in our understanding of how AAC(6′)-Ib mediates resistance is its intracellular dynamics. Previous studies have demonstrated that gene amplification is correlated with higher expression of aminoglycoside modifying enzymes and increased resistance levels (23–29). However, these studies rely on the number of gene copies rather than the correlation between the measured number of protein molecules and resistance levels. In this work, we describe the design of a fluorescence-based method to accurately determine the number of molecules of AAC(6′)-Ib and a CRISPR interference (CRISPRi) system to regulate the number of AAC(6′)-Ib molecules synthesized. We observed a correlation between the quantity of AAC(6′)-Ib molecules and amikacin resistance up to a threshold where adding additional AAC(6′)-Ib molecules ceases to be associated with increased resistance. Live cell imaging showed that AAC(6′)-Ib aggregates at the poles at higher concentrations, a process that may denature and inactivate the excess molecules. Single molecule microscopy of the dynamics of AAC(6′)-Ib in the cellular context showed that the enzyme freely diffuses in the cytoplasm, not binding to any particular cellular structure. Our results provide a quantitative understanding of the relationship between the number of enzyme molecules and resistance levels. Furthermore, our findings suggest that the upper limit to amikacin resistance arises due to protein aggregation at increased intracellular concentration.

## Results

### Mutant gRNAs mediate progressive regulation of AAC(6′)-Ib gene expression

To fine-tune AAC(6′)-Ib expression levels, we used CRISPRi. This method relies on a catalytically dead variant of Cas9 (dCas9) that binds a specific DNA sequence but fails to cleave it, producing a roadblock for transcriptional initiation or elongation (30). The level of repression of *aac(6′)-Ib* expression is maximum if the guide RNA (gRNA) and the targeted DNA sequence are perfectly complementary. Therefore, introducing mismatches in the gRNA reduces the repression strength (31, 32). We generated a library of gRNAs that permitted the expression of different quantities of AAC(6′)-Ib molecules, which were quantified using a derivative of the pJHCMW1 plasmid that carries the native *aac(6′)-Ib* gene fused to the gene coding for the mNeonGreen fluorescent protein (Fig. 1A) (14). We introduced this plasmid into *E. coli* strains harboring an IPTG-inducible dCas9-coding gene and a constitutive gRNA-coding gene (Fig. 1B). The gRNA molecules in each strain were designed to target different *aac(6′)-Ib* regions (the promoter, the coding sequence, or the linker, the coding or non-coding strand), and to have various degrees of complementarity to the target sequence (Fig. 1A and Fig. S1). The effect of each gRNA on the AAC(6′)-Ib expression level was assessed by spectrofluorometry on overnight LB cultures supplemented with IPTG (Fig. 1C). Correcting the total intensity by the culture’s optical density and the corresponding colony-forming units allows an estimation of the protein’s copy number per cell. This spectrofluorometry-based quantification of AAC(6′)-Ib molecules per cell was corroborated by confocal microscopy (Fig. S2-3). The results confirmed that dCas9 blocks the initiation and elongation of transcription and that the length of complementarity to the target sequence is correlated with the strength of repression. Targeting either strand of the *aac(6′)-Ib* promoter provides strong transcriptional repression (Fig. 1D). Guiding dCas9 binding to the coding strand of the structural gene or the linker results in robust inhibition of gene expression. Conversely, repression is not as pronounced when targeting the non-coding strand (Fig. 1E and F). Reduction of the length of complementarity to the coding strand was associated with a decrease in the degree of inhibition of gene expression (Fig. 1E and F). The results described in this section indicate that the fluorescence-fusion system is adequate to assess the number of AAC(6′)-Ib molecules per cell, and the CRISPRi fluorescence-based system is an efficient tool to modulate the AAC(6′)-Ib copy number, spanning three orders of magnitude.

**Figure 1.**
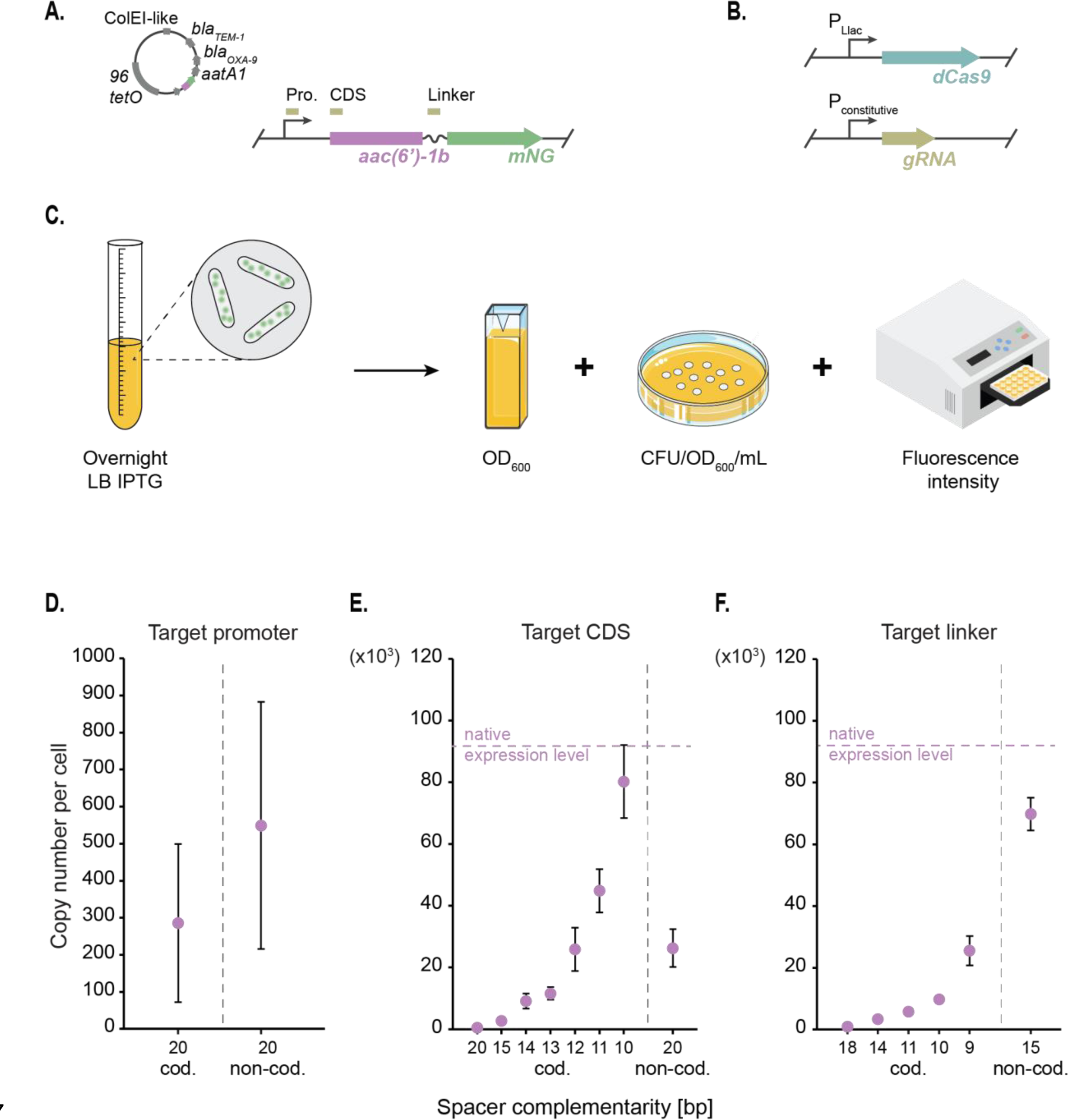
– Regulation of AAC(6′)-Ib copy number by CRISPRi. **A.** Schematic representation of the pJHCMW1 plasmid and a zoomed view on the gene of interest, the aminoglycoside acetyltransferase AAC(6′)-Ib tagged with mNeonGreen (mNG) under its native promoter. Boxes above the gene mark the regions targeted by the dCas9, which include the promoter (Pro.), coding sequence (CDS), and linker. **B.** Schematic representation of the dCas9-coding gene under the PLlac promoter and the guide RNA (gRNA) with its constitutive promoter. **C.** Illustration of the workflow for AAC(6’)-Ib protein copy number quantification. The copy number per cell is estimated based on the optical density (OD600), bacterial count (colony forming units (CFU)/OD600/mL), and intensity measured with a spectrofluorometer from an overnight LB culture of the strain that carries the pJHCMW1 derivative plasmid and expresses the dCas9 protein. **D.** Average AAC(6′)-Ib copy number per cell obtained with the spectrofluorometer for strains harboring gRNAs with different complementarity to either the coding (cod.) or the non-coding strand (non-cod.), varying lengths and targeting the promoter sequence, the CDS (**E.**) or the linker (**F.**) of the gene. The native expression level represents AAC(6’)-Ib copy number per cell in the absence of dCas9 repression. Error bars represent the standard deviation.

### At low copy numbers, the abundance of AAC(6′)-Ib and the minimum inhibitory concentration of amikacin follow a linear relationship

We carried out microtiter plate-based assays to evaluate how variations in the cellular copy number of AAC(6′)-Ib impact amikacin-resistance levels. We determined the amikacin concentration needed to reduce the culture OD_600_ value by 50%, defined as the inhibitory concentration or IC_50_. The native copy number of AAC(6′)-Ib in the strains used is ∽92,000 molecules per cell, and the corresponding measured IC_50_ of amikacin was 135 µg/mL (Fig. 2A). Although such a high copy number is surprising, it is congruent with expression of the gene from 20-30 copies of the plasmid from a strong promoter (14, 17). Within a range spanning from none to about 10,000 AAC(6′)-Ib copies per cell, the IC_50_ value increased as a function of the copy number, with an apparent relation of 1 µg/mL resistance increase for every 170 AAC(6′)-Ib copies per cell (Fig. 2B). However, it was of interest that this correlation ceased at ∽10,000 – 20,000 copies per cell when the resistance level hit a plateau at IC_50_ of amikacin of 105 µg/mL. This value was constant, up to ∽70,000 copies per cell, where the IC_50_ increased again up to 135 µg/mL (Fig. 2A). Note that this plateau does not seem to arise from the mNeonGreen tag as it does not impair the activity of the enzyme (Fig. S4).

**Figure 2.**
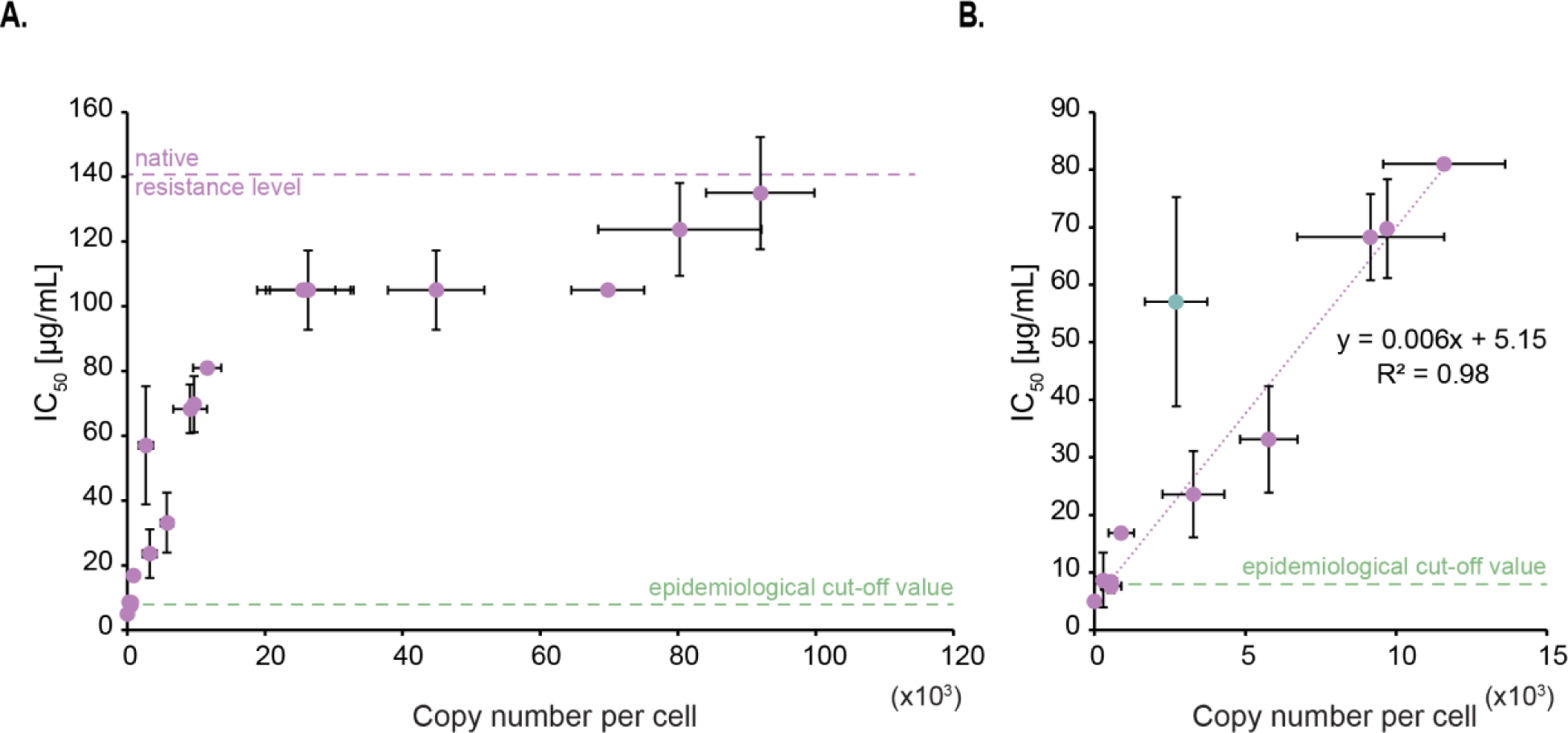
– Impact per AAC(6′)-Ib enzyme molecule on amikacin resistance level. **A.** Correlation between the inhibitory concentration at which 50% of the maximal culture density is observed (IC50) and AAC(6′)-Ib copy number per cell. The native resistance level of the wild-type plasmid and the epidemiological cut-off value are also indicated (https://www.eucast.org). Error bars represent standard deviations. **B.** Lower copy numbers follow a linear relation with IC50 to amikacin. The linear trendline equation and its corresponding R-squared value are shown. The green dot represents an outlier that is excluded from the trendline.

As a point of reference, the “epidemiological cut-off value,” which is defined as the highest minimum inhibitory concentration (MIC) of the wild-type population, is reported to be 8 µg/mL for enterobacteria (http://www.eucast.org) (33). In our conditions, a concentration of 8 µg/mL is reached when the copy number of AAC(6′)-Ib per cell is ∽475. While the conditions used to estimate this epidemiological cut-off value differ from those used in this work, we expect that the minimum copy number of the enzyme needed to be classified as resistant to amikacin should at least be in the same order of magnitude as our estimate.

### AAC(6′)-Ib forms aggregates at higher concentrations

To gain insight into the correlation between protein copy number and resistance level described in the previous section, we further assessed the number of molecules per cell using confocal microscopy. The results of these assays showed that both techniques, spectrophotometry and microscopy, produced numbers per cell within the same order of magnitude (Fig. S3). However, it was surprising to us that microscopy permitted us to observe that the enzyme distributes homogeneously in the cytoplasm up to a specific concentration, after which any increase tends to result in the formation of aggregates at the cell poles (Fig. 3A). The mNeonGreen fluorescent protein does not tend to aggregate at very high concentrations, suggesting that appearance of these inclusion bodies are independent of the presence of the fluorescent tag (Fig. S4). Low solubility and tendency to aggregate as inclusion bodies have also been observed in a related AAC(6’)-Ib variant (18).

**Figure 3.**
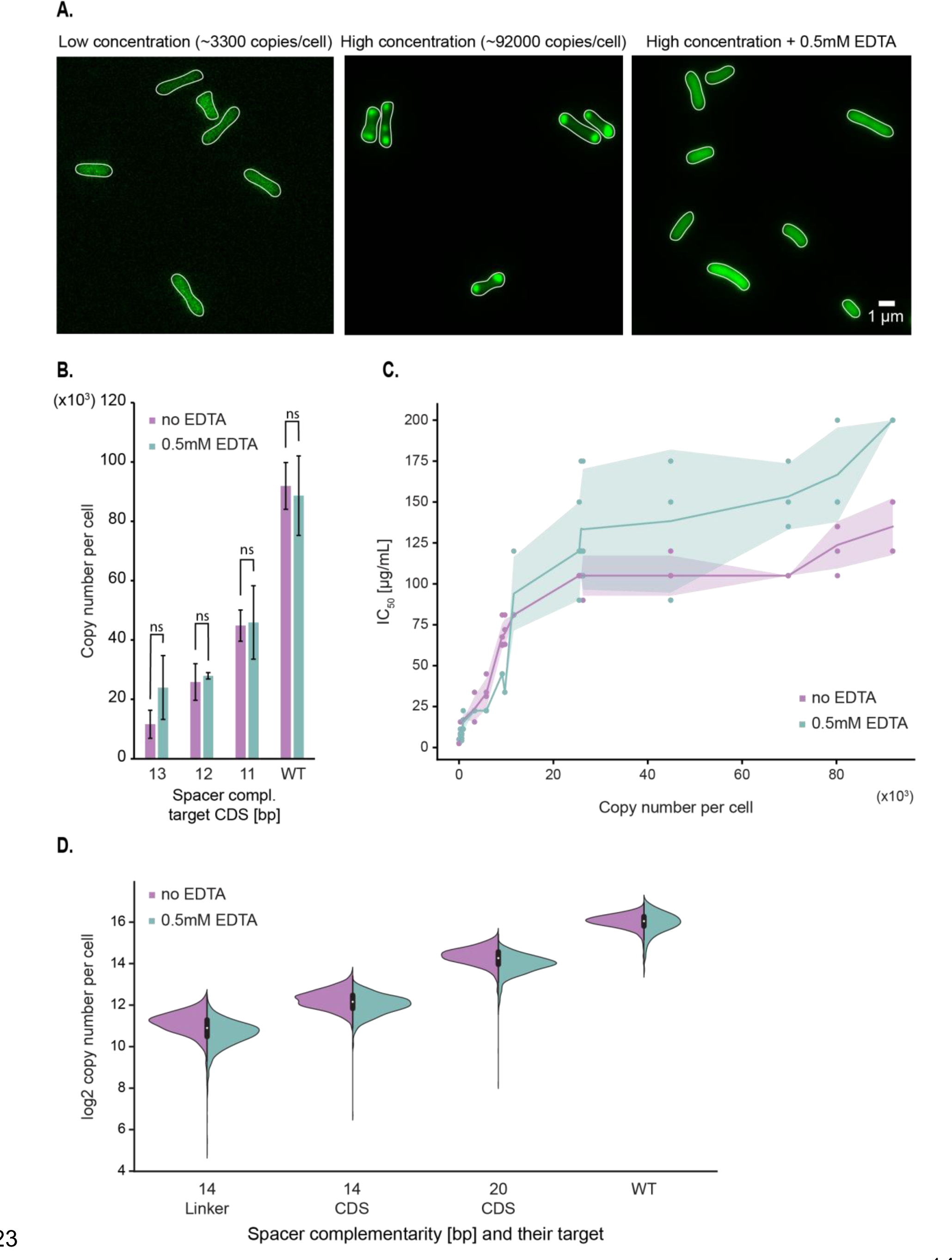
– Formation of AAC(6′)-Ib aggregates at high protein concentrations partially inhibit enzyme activity against amikacin. **A.** Representative images of strains carrying the pJHCMW1 plasmid derivative that encodes the *aac(6′)-Ib-mNeonGreen* gene and the CRISPR-dCas9 system that targets the coding sequence (CDS) of this gene, with a strong repression on the left and no repression in the middle image. The rightmost image displays the wild-type strain in presence of 0.5 mM EDTA (ethylenediaminetetraacetic acid). **B.** Average copy number per cell of AAC(6′)-Ib in the presence or absence of 0.5 mM EDTA, measured by spectrofluorometry. Each strain carries different lengths of gRNA complementarity to the CDS of the *aac(6′)-Ib* gene. Error bars represent the standard deviation. A statistical analysis compared the two conditions (ns, not significant for p=0.05). **C.** Scatterplot showing the relationship between AAC(6′)-Ib copy number per cell and IC50 to amikacin. The shaded area represents the standard deviation in IC50. **D.** Distribution of AAC(6′)-Ib copy number, estimated with the confocal microscope, for strains encoding different guide RNAs in the presence or absence of 0.5 mM EDTA. Complementarity of the gRNA to the coding strand of the *aac(6′)-Ib* gene is indicated for each strain.

The observed plateau in amikacin resistance may arise from AAC(6′)-Ib’s inability to exist in soluble and native form at copy numbers higher than 10,000 – 20,000. To test this hypothesis, cells were cultured in broth supplemented with the chelating agent ethylenediaminetetraacetic acid (EDTA), a compound known for permeabilizing bacterial outer membranes, disrupting ionic interactions and favoring solubilization of protein aggregates (34, 35). Fig. 3A shows that the fluorescent protein was homogeneously distributed over the cytoplasm in cells cultured in the presence of EDTA. In addition, Fig. 3B shows that the addition of EDTA did not modify the number of molecules per cell, a clear indication that the sole effect of EDTA was to dissolve the polar aggregates. Furthermore, the amikacin IC_50_ for cells cultured in broth supplemented with EDTA increased as the AAC(6′)-Ib copy number increased, but the new distribution still contained a plateau (Fig. 3C). A comparison of the amikacin resistance levels in cells producing a number of molecules consistent with the formation of aggregates but that were cultured in the presence of the chelator showed a 1.5-fold increase in resistance. We conclude that aggregation of AAC(6′)-Ib must inhibit enzymatic activity and is, at least in part, responsible for the observed plateau effect.

### Diversity of AAC(6′)-Ib expression profile per cell within a population

While estimations of protein expression per cell are reproducible, measurements made with the fluorometer do not provide any information regarding the variability of gene expression within a population. The confocal microscope, however, allows an estimation of AAC(6′)-Ib copy number at the single-cell level, revealing potential cell-to-cell variation. This is particularly important in the case of strains where the expression of AAC(6′)-Ib is regulated by CRISPRi. If there was cell-to-cell variability in the levels of dCas9, the individual cell’s AAC(6′)-Ib copy numbers could be significantly different, which would cloud the interpretation of our results. However, the results showed that AAC(6′)-Ib expression is normally distributed in the cell population of individual strains (Fig. 3D and Table S4). Also, cell-to-cell fluctuations in the protein concentration were similar to changes at native expression levels. These results indicated that our CRISPRi system has no impact on the intrinsic variations of AAC(6′)-Ib expression. Importantly, this experiment also confirmed that the presence of EDTA in the cell’s environment does not modify the protein’s copy number or cell-to-cell variability.

### AAC(6′)-Ib diffuses freely in the cell

To further understand the dynamics of AAC(6′)-Ib within the cytosol, we characterized its diffusion properties using single-particle tracking via Photoactivated Localization Microscopy (sptPALM) (36). We used a fusion of AAC(6′)-Ib with the photoconvertible fluorescent protein mMaple (37) and captured images at 20-ms rates. To study the dynamics of single molecules, we exposed the whole field of view to continuous 405 nm light at low intensity, conditions that result in the activation of a single molecule of mMaple per cell, on average. The resulting images permitted us to track a molecule’s position and follow its trajectory (Fig. 4A). Tracked molecules of AAC(6′)-Ib covered all the cytosol, with no location preference or obvious binding to any cellular structure. The average apparent diffusion coefficient of AAC(6′)-Ib diffusing in the cell is the same as that of mMaple alone (Fig. 4B). In conclusion, the results described in this section indicate that in the absence of aggregation, AAC(6′)-Ib molecules diffuse freely across the cytosol.

**Figure 4.**
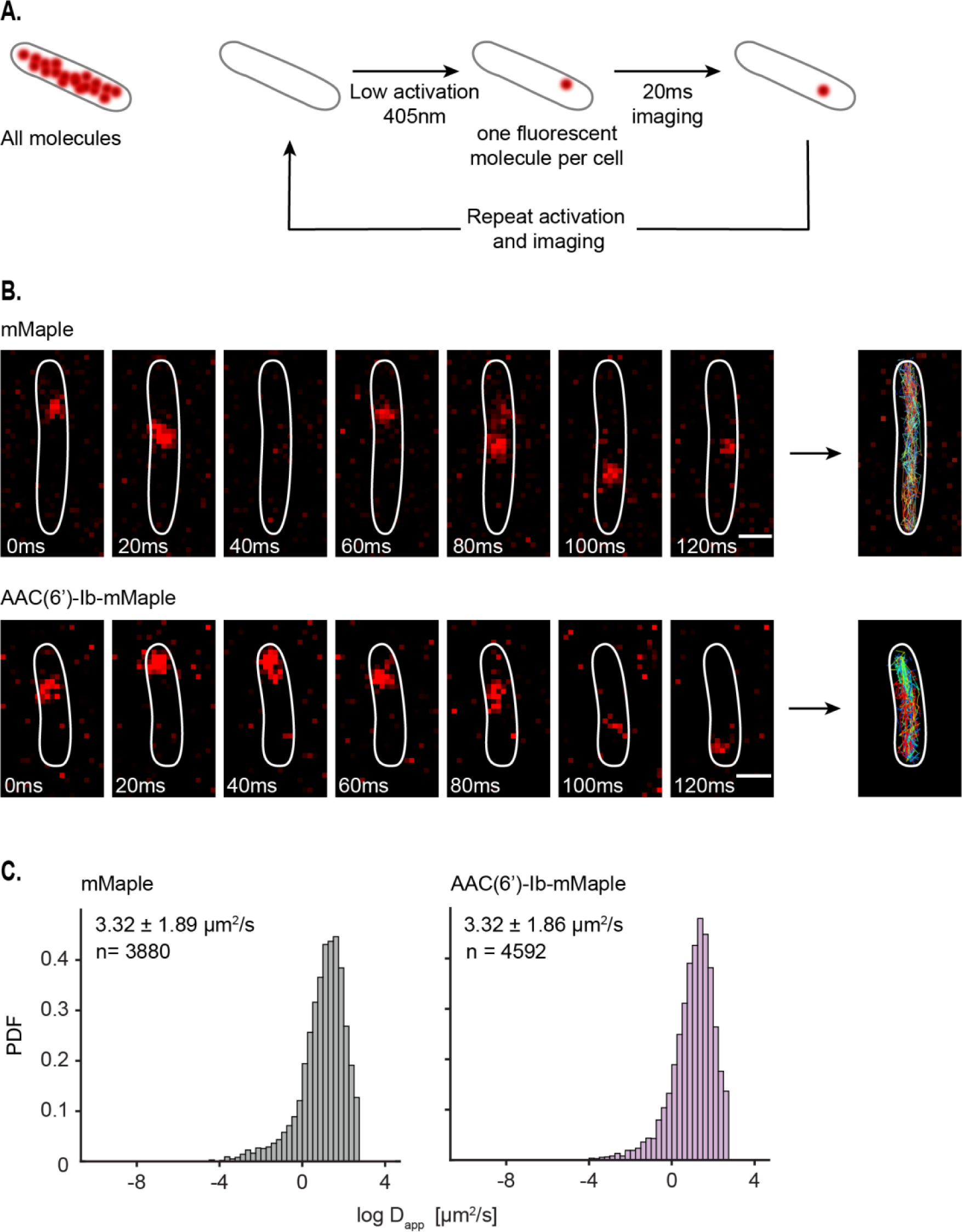
– AAC(6′)-Ib is highly dynamic in the cells. **A.** Principle of single-particle tracking via Photoactivated Localization Microscopy (sptPALM). **B.** Representative images of mMaple and AAC(6′)-Ib-mMaple foci every 20 ms. Tracks are represented on the rightmost images. Scale bar: 1µm. **C.** Distribution of the logarithm of the apparent diffusion coefficient (Dapp) for mMaple (left) and AAC(6′)-Ib-mMaple (right). The y axis represents the probability density function (PDF). The apparent diffusion coefficient of the diffusing fraction, taken from the exponential of the mu and sigma obtained from the Gaussian distribution, and the total amount of tracks analyzed are indicated.

## Discussion

The rapid dissemination of aminoglycoside modifying enzymes has severely reduced the effectiveness of aminoglycosides. The acetyltransferase AAC(6′)-Ib is among the most clinically relevant due to its ability to catalyze the inactivation of numerous aminoglycosides used to treat severe infections. The substrates of AAC(6′)-Ib include natural as well as semisynthetic aminoglycosides such as amikacin (7, 11, 13). The *aac(6′)-Ib* gene, as well as the AAC(6′)-Ib protein, have been the subject of intense genetic, structural, and biochemical studies (7, 13, 18, 19, 38, 39). However, despite these advances, numerous unanswered questions remain; clarifying these is critical for the rational design of novel therapies.

To reduce the knowledge gap, we studied the dynamics of the AAC(6′)-Ib protein inside the cytosol to assess how its copy number variation impacts amikacin resistance. To do so, we devised methodologies to quantify and control AAC(6′)-Ib copy number. A fusion AAC(6′)-Ib-mNeonGreen enabled us to quantify the number of protein molecules per cell using fluorometry and confocal microscopy. Then, we used a CRISPRi system to produce strains that expressed a wide range of molecules per cell. These tools were used to compare amikacin resistance levels in strains harboring different numbers of enzyme molecules. We observed a close correlation between resistance levels and the number of protein molecules at low concentrations. These results agree with previous reports that the copy number of different resistance genes is amplified in mutants displaying increased resistance (23–29). However, previous studies were limited to correlating the number of gene copies, increased by amplification mechanisms or modification of plasmid copy numbers and resistance levels. Absent in these reports are correlations of actual expressed resistance enzyme molecules and resistance levels. This work now accurately measured amikacin resistance as a function of AAC(6′)-Ib copy number.

Our results indicate that resistance levels reach a plateau at a certain level of AAC(6′)-Ib concentration. The single-cell results obtained by microscopy showed that the AAC(6′)-Ib molecules tend to aggregate and localize at the poles in structures resembling inclusion bodies at higher concentrations. This observation, taken together with the well-known fact that the protein has limited water solubility (18), explains, at least in part, the lack of correlation between the AAC(6′)-Ib copy number and resistance levels at higher protein concentrations. Physical exclusion of AAC(6′)-Ib, due to the formation of inclusion bodies, can limit its accessibility to the antibiotic molecules. Adding EDTA to the growth medium quantified the contribution of aggregate formation to the observed plateau in resistance levels. The chelator’s presence interfered with the formation of AAC(6′)-Ib aggregates and was correlated with an increase in the amikacin IC_50_. Not only do these results associate aggregation with increased resistance, but they also suggest that the process is structurally dynamic and reversible. This behavior will be essential to consider when developing strategies to inhibit resistance. For example, antisense inhibition of AAC(6′)-Ib expression, a methodology that attempts phenotypic conversion to susceptibility by turning off gene expression, will need to be very efficient to reduce the number of proteins produced well below the aggregation threshold.

Another potential contributor to the plateau observed is enzyme kinetics. The classic expectation that the enzymatic reaction velocity increases linearly with enzyme concentration is valid only when the substrate is present at much higher concentrations than the enzyme. For amikacin resistance, substrates inside the cytosol may not saturate the enzyme. The intracellular concentrations of acetyl-CoA, the acyl donor, have been reported to be in the 20-600 μM range (40), which are comparable to those estimated for AAC(6′)-Ib in strains producing high copy numbers (40-150 μM). Availability of acetyl-CoA may limit the enzyme activity rate. Also, amikacin concentrations inside the cytosol are likely lower than those in the growth medium (note that 125 μg/ml equals 160 μM). Hence, the substrates/enzyme concentration ratios may also influence the resistance levels in growing cells. While the enzyme solubility issue seems to affect our results, future experiments must be planned to help understand the enzyme kinetics in the cytosol. Adding to these complications, one must consider that different bacteria will have different permeabilities to the antibiotic and could possess efflux pumps.

Our previous studies showed that unlike other resistance enzymes like β-lactamases that are periplasmic (41), AAC(6′)-Ib is uniformly distributed within the cytoplasm (16, 42). However, those experiments required overexpression of the enzyme, which could have modified the subcellular location at physiological levels. The methodology used in this work corroborated our early results and further illustrated the dynamics of AAC(6′)-Ib inside the cytosol. The protein can traverse the length of the cell in ∼100 ms.

In conclusion, this work makes inroads in understanding the individual contribution of AAC(6′)-Ib molecules to resistance to aminoglycosides and clarifies its dynamics inside the cell. Importantly, it opens the doors to understanding the role of solubility and expression levels in the resistance phenotype. We expect that the system we devised will help us better understand antibiotic resistance from cellular and molecular biology perspectives.

## Materials and methods

### Strains and growth conditions

All strains used are derivatives of AB1157 (Table S1). Cells were routinely grown in LB or M9 minimal media. M9 was supplemented with glycerol (final concentration 0.2%); 100 µg/ml of amino acids threonine, leucine, proline, histidine, and arginine; and thiamine (0.5 µg/ml). When required, antibiotics were added at the following concentrations: ampicillin (100 µg/ml), kanamycin (50 µg/ml), and chloramphenicol (25 µg/ml). The TB25 strain was constructed by chromosomal integration of the *P_lacq_-lacI P_Llac-s_-dCas9* gene from the pTB35 plasmid at the lambda *attB* site (43). Each gRNA was inserted into the pTB40-1 plasmid through site-directed mutagenesis that was carried out by uracil-specific excision reagent (USER) cloning (Table S2) (44). The different gRNAs were further integrated by lambda red in the *argE* gene. OD030 was constructed using lambda red recombination with the pROD93 plasmid carrying the *mMaple* gene, which was inserted into the *galK* gene. The mNeonGreen gene was integrated into the pTT4 plasmid (17) to generate pTT4-mNG-wL, and the construction was subsequently sequenced by whole plasmid sequencing service of plasmidsaurus (www.plasmidsaurus.com). The pFH3 plasmid is a pTT4-mNG-wL derivative where the *mNeonGreen* gene was replaced by *mMaple3* by *Nco*I digestion and ligation. All primers used for these constructions are described in Table S3.

### Protein expression and purification

Fluorescent proteins were produced using pROD93 expression vector in BL21(DE3) strains. Cells were grown at 37°C until they reached an OD_600_ ∼ 0.6. Protein expression was then induced by adding 0.2% of L-arabinose and incubating the culture at 30°C. After 4 hours of induction, bacteria were centrifuged and the pellet was resuspended in Resuspension Buffer (25 mM Tris HCl pH 7.5, 250 mM NaCl) supplemented with one tablet of cOmplete protease inhibitor (Roche, Cat # 04693159001). Cells were then lyzed using a high-pressure Emulsiflex C5 homogenizer (Avestin), and the lysate was cleared by ultra-centrifugation (35 000 rpm, 1 hour, 4°C). The lysate was added to the Ni-IDA resin (Takara, Cat # 635660) equilibrated with Binding Buffer (50 mM NaPi, 20 mM Imidazole, 300 mM NaCl, 10% glycerol, pH 8) and incubated for at least an hour at 4°C. The resin was transferred to a disposable column and washed out at least 5 times with Binding Buffer and twice with Binding Buffer that has a higher concentration of imidazole (40 mM). Protein was eluted in Elution Buffer (50 mM NaPi, 200 mM Imidazole, 300 mM NaCl, 10% glycerol, pH 8). The protein was then dialyzed in Slide-A-Lyzer MINI Dialysis Devices immersed in the Storage Buffer (25 mM Tris HCl pH8, 300 mM NaCl, 10% glycerol) and finally stored, protected from light, at -20°C.

### Fluorometer calibration assay

For mNG fluorescence calibration, the purified protein stock was first diluted in M9 minimal media to a concentration of 2.5 10^-2^ mg/mL and then diluted 1:2 for a total of 7 dilutions. 200µL of each concentration was distributed in a black microplate (Fisher, Cat # 7200590), and the total intensity was measured with a Cary Eclipse Fluorescence Spectrophotometer (Agilent Technologies) at 500 nm for excitation and 530 nm for emission. The slits were set to 10 and 20 nm for excitation and emission, respectively. After correcting for the background fluorescence of M9 minimal media, we could estimate the average fluorescence intensity per mNG protein.

### Copy number quantification using the spectrofluorometer

The strains expressing the dCas9 with the different gRNAs and carrying the pTT4-mNG-wL plasmid were first grown overnight in LB media in the presence of ampicillin and 1 mM IPTG at 37°C. The OD_600_ of the culture was measured, and 1 mL was washed in M9 minimal media, from which 200 µL were distributed in a clear 96-well plate as triplicates. Using the same parameters as in the fluoromoter calibration assay, the total fluorescence was measured using the Cary Eclipse fluorometer. AAC(6′)-Ib copy number could thus be estimated by dividing the measured intensity by the OD_600_, correcting for the fluorescence of the cells without the fluorescent plasmid and dividing by the average fluorescence intensity per mNG molecule (4.92 x 10^-12^, Fig. S2). This value was then divided by the estimated bacterial count (5.22 x 10^8^ CFU/OD_600_/mL) to calculate the copy number per cell.

### Copy number estimation with the confocal microscope

The AAC(6’)-Ib copy number per cell was confirmed using the confocal microscope. Single-molecule mNG intensity was first quantified using the YHZ23 strain that carries a mNG-tagged Nup59, a well-characterized 16-mer component of the nucleopore complex (NPC) in budding yeast. This strain was imaged using a custom-built spinning disk confocal microscope (Leica DMi8 inverted microscope with Quorum Diskovery platform, 50 µm pinhole spinning disk and two iXon Ultra 512×512 EMCCD cameras). A single colony was grown in synthetic complete (SC) media, with shaking, for 5 hours at 30°C. This culture was diluted 100 times and cultured overnight in the same conditions before being diluted again to an OD_600_ of 0.15 and incubated until it reached 0.3. 1mL of this culture was spun down and concentrated twice in fresh SC media before being spread on a 1% agarose pad. Imaging was performed by taking 10 z-stacks of 0.59µm step size with 488 nm laser at 25% intensity and with a 200ms exposure time. Single spots were identified manually throughout the z-stacks and were subsequently fitted to 2D Gaussian with the GaussFit on Spot plugin in ImageJ. Using Matlab, the intensity values were fitted to a Gaussian Mixture Model, resulting in two distinct components (Nup59 from single and double NPCs). The mean of the first component, *i.e.,* in the single NPC, is the intensity of 16 mNG molecules (mean intensity = 11374.4 and thus intensity of a single mNG = 710.9).

As for the fluorometer-based quantification, bacterial cultures were incubated overnight in LB media in the presence of ampicillin and 1 mM IPTG at 37°C. Imaging was performed by taking 7 z-stacks of 0.59µm step size with 488 nm laser at 25% intensity and with a 200ms exposure time. Stacked images were projected for analysis by summing all pixel values at each position using Fiji. Images were then segmented with Cellpose (45), and the total intensity per cell was quantified using custom Fiji and Python scripts. In short, the Mean value (total intensity in the cell divided by its area) was corrected for the average Mean value in cells that do not carry the fluorescent protein, divided by the intensity per mNG molecule estimated for Nup59-mNG and finally corrected by the average cell area.

### IC_50_ determination

Cells were first cultured overnight in LB medium in the presence of ampicillin and 1 mM IPTG. They were then diluted 1:10000 in fresh LB, and 5µL was added into each well of a 96-well microtitre plate (Sarstedt, Cat # 82.1581.001) carrying 100µL LB of a series of 2-fold dilutions of amikacin as well as 1mM IPTG. When desired, the LB media was also supplemented with 0.5 mM EDTA. The plate was then incubated for 20 hours at 37°C. IC_50_ was defined as the lowest antibiotic concentration at which half of the bacterial growth is inhibited.

### Single molecule experiments and analysis

Overnight bacterial cultures were diluted in M9 minimal media supplemented with 1 mM IPTG and cultured until they reached early logarithmic phase (OD_600_ of 0.1-0.2). Cells were then visualized with an inverted Olympus IX83 microscope at room temperature. A Hamamatsu Orca-Flash 4.0 sCMOS camera was used for image capturing, and Z-stacks were done using a NanoScanZ piezo by Prior Scientific. Excitation was performed with an iChrome Multi-Laser Engine from Toptica Photonics, combined with a 405/488/561/640nm filter set (Chroma). We used a Single-line cellTIRF illuminator (Olympus) for the experiments, and the Olympus CellSens 2.1 imaging software was used to control the microscope and the lasers. All data analyses were performed using custom MATLAB scripts (Mathworks), except for single molecule tracking that was done using Trackmate (46, 47).

### AAC(6’)-Ib enzymatic activity assay

Total soluble proteins (enzymatic extracts) were prepared as before (48). Briefly, cells were pelleted from cultures by centrifugation and resuspended in a 0.5 mM MgCl2 solution. The cells were lysed by sonication with a Heat Systems Ultra-sonic, Inc., Model No. H-IA (Plainview, NY, USA) cell disrupter. The soluble protein fraction was then separated from unbroken cells, membranes, and cell debris by centrifugation in a microfuge for 10 min at 4 °C. The protein concentration of the extracts was measured using a commercial reagent (Bio-Rad Protein Assay). Acetyltransferase activity was assessed using the phosphocellulose paper binding assay [Haas & Dowding, 1975]. Soluble extract (120 μg protein) obtained from E. coli TOP10(pUC57AAC2Ia) cells was added to the reaction mixture (200 mM Tris HCl pH 7.6 buffer, 0.25 mM MgCl2, 330 μM plazomicin, the indicated concentrations of sodium acetate or silver acetate, and 0.05 μCi of [acetyl-1-^14^C]-acetyl-coenzyme A (specific activity 60 μCi/μmol). The reaction mixture final volume was 30 μL. Silver ions were added as silver acetate due to its adequate solubility in water. After incubating the reaction mixture at 37 °C for 30 min, 20 μL were spotted on phosphocellulose paper strips. The unreacted radioactive substrate [acetyl-1-^14^C]-acetyl-coenzyme A was removed from the phosphocellulose paper strips by submersion in 80 °C water followed by two washes by submersion in room temperature water. After this treatment, the only radioactive compound bound to the phosphocellulose paper strips was the acetylated plazomicin. The phosphocellulose paper strips were then dried and the radioactivity corresponding to enzymatic reaction product was determined in a scintillation counter.

## Acknowledgments

We thank Jia Yin Xiao for sharing microscope single-molecule calibration data. The Reyes lab was funded by the Natural Sciences and Engineering Research Council of Canada (NSERC RGPIN-2019-05701), the Canadian Institutes for Health Research (CIHR PJT-162247), the Canada Foundation for Innovation (CFI# 228994), and the Canada Research Chairs program. The Tolmasky lab was funded by Public Health Service Grant 2R15AI047115-06 (National Institute of Allergy and Infectious Diseases, NIH) and the Ramirez lab was funded by Public Health Service Grant SC3GM125556 (National Institute of General Medical Sciences, NIH). Fong Hue was supported by grant MHRT 2T37MD001368 from the National Institute on Minority Health and Health Disparities, NIH. Some of the experiments used equipment from the Integrated Quantitative Bioscience Initiative (IQBI), funded by CFI 9.

## Author contributions

O.D.D.: Conceptualization, Formal analysis, Investigation, and Writing. F.H.: Formal analysis and Investigation, T.Y: Formal analysis and Investigation. L.W.: Formal analysis and Investigation. M.L.: Formal analysis and Investigation. L.R.: Formal analysis and Investigation. T.T.: Formal analysis and Investigation. K.P.: Formal analysis and Investigation. M.S.R.: Formal analysis and Investigation. W.R: Conceptualization and Writing. M.E.T.: Conceptualization, Supervision, Funding acquisition and Writing. R.R.-L.: Conceptualization, Supervision, Funding acquisition and Writing.

## Supplementary data

**Figure S1.**
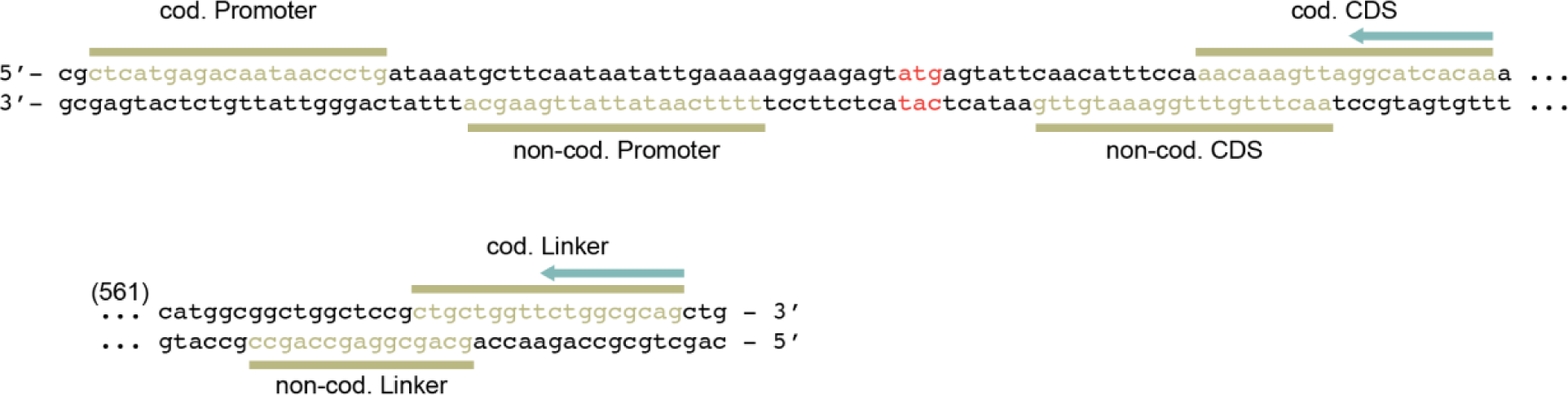
– dCas9 targets (Promoter, CDS and Linker on both the coding (cod.) and non-coding (non-cod.) strands) in the *aac(6′)-Ib* gene on the pJHCMW1-derivative plasmid. The complementary sequence to the gRNA (green bar) is highlighted in green and the blue arrow represents the positions at which the mismatches in the sequence of the gRNA were introduced. The *aac(6′)-Ib* START codon is indicated in red and the number of base pairs not included in this representation are indicated in parenthesis.

**Figure S2.**
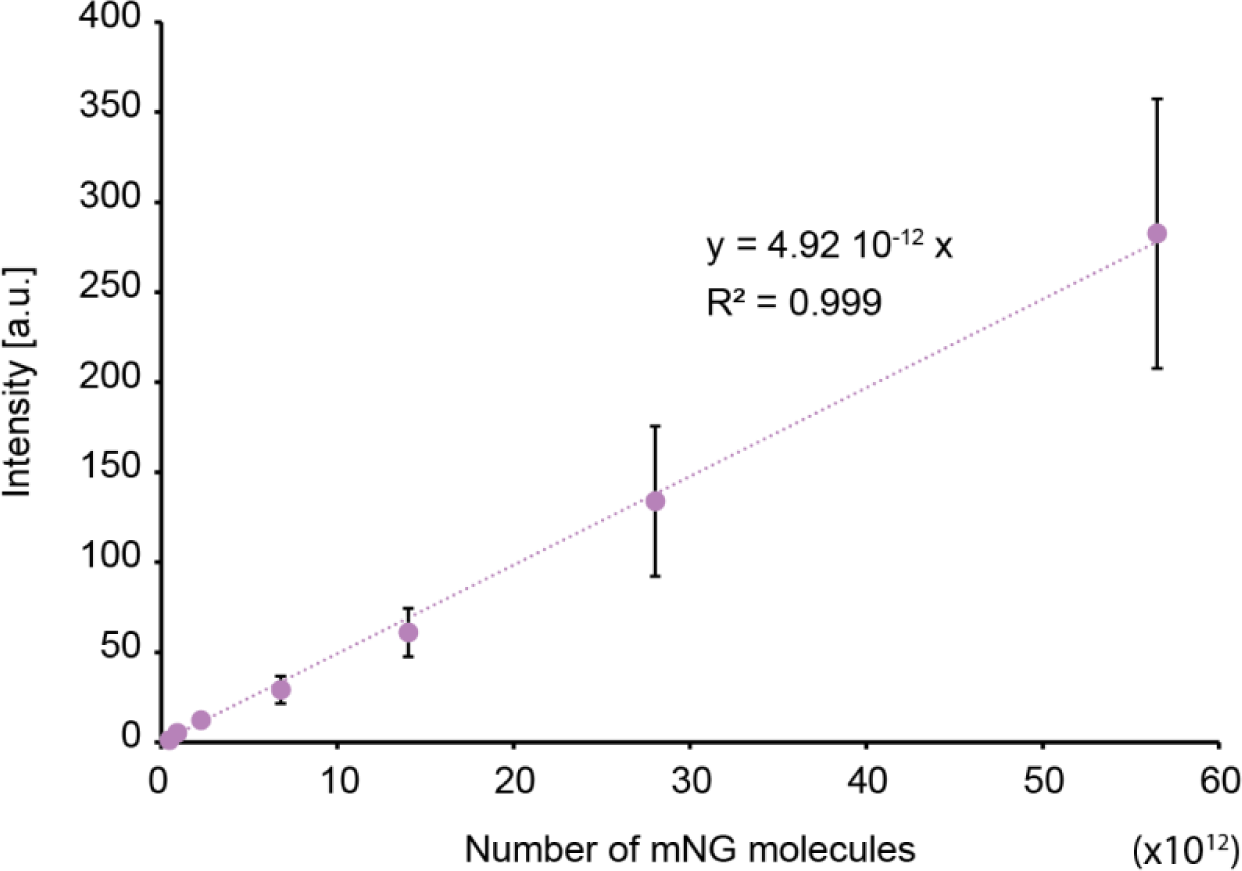
– Calibration curve between mNeonGreen intensity measured by spectrofluorometry (in arbitrary units, a.u.) and the number of mNeonGreen molecules. The equation of the linear trendline, as well as the correlation coefficient (R^2^), are indicated on the plot.

**Figure S3.**
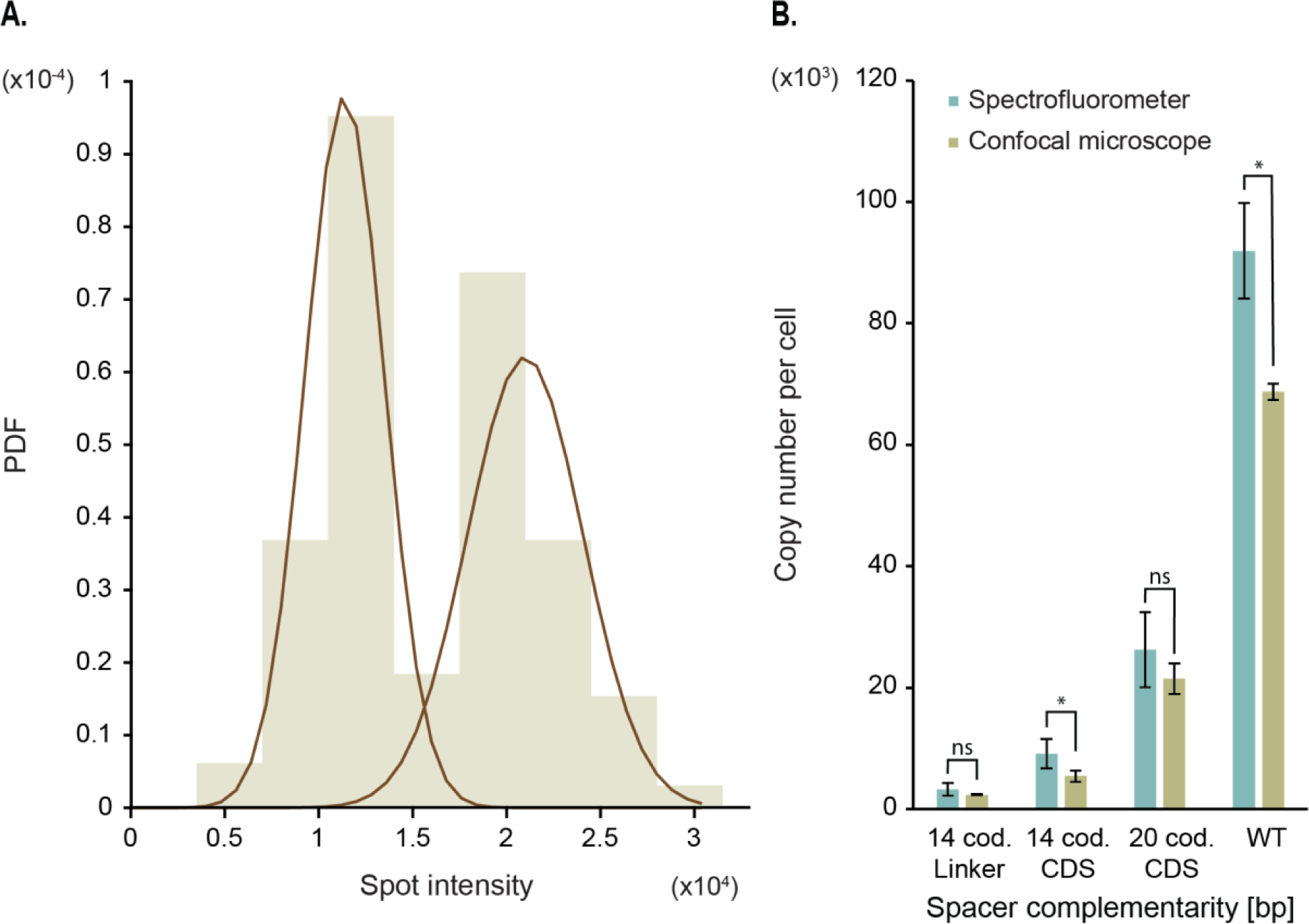
– Confirmation of AAC(6′)-Ib copy number by confocal microscopy. **A.** mNeonGreen intensity per molecule, estimated by imaging Nup59-mNeonGreen. The brown lines represent the Gaussian Mixture Model fitted to each population. The one on the left represents the 16-mer Nup59 (mean intensity = 11374.4), and the second peak carries 32 Nup59 (mean intensity = 20988.6). This gives an average intensity of 710.9 per mNeonGreen molecule. In total, 93 spots are represented in this plot. **B.** AAC(6′)-Ib copy number estimated with the spectrofluorometer (blue bars) and with the confocal microscope (green bars) for four strains, each carrying a different guide RNA. The length of the gRNA that is complementary to the coding (cod.) strand, as well as the targeted region of the *aac(6′)-Ib* gene, are indicated. Error bars represent the standard deviation and comparative statistical analysis was performed to compare the two methods (ns, not significant, and *, significant for p=0.05).

**Figure S4.**
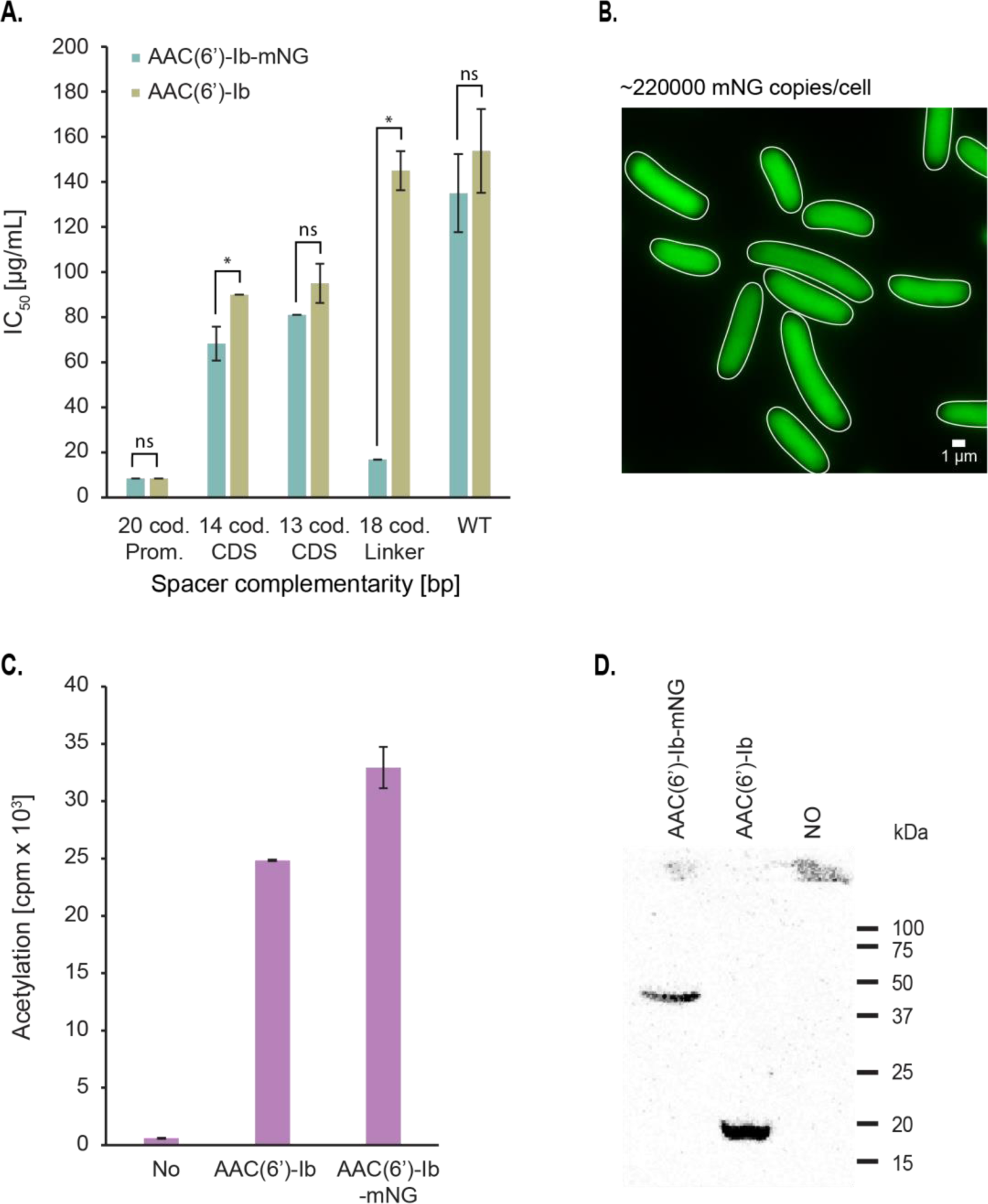
– The mNeonGreen (mNG) fluorescent tag does not alter AAC(6’)-Ib activity and does not force protein aggregation. **A.** Inhibitory concentration at which 50% of growth is inhibited by the amikacin antibiotic (IC50) for different repression levels of AAC(6’)-Ib copy number per cell. The length of the gRNA that is complementary to the coding (cod.) strand, as well as the targeted region of the *aac(6′)-Ib* gene, are indicated (Prom, promoter; CDS, coding sequence; WT, wild-type). Error bars represent the standard deviation and comparative statistical analysis was performed to compare the effect of the mNG tag on the activity of the enzyme in the cell (ns, not significant, and *, significant for p=0.05). **B.** Representative image of AB1157 cells expressing high concentrations of mNG molecules from a pUC18 derivative plasmid. Copy number quantified by spectrofluorometry (221483.5 ± 16440.8 mNG copies/cell). **C.** Effect of mNG on AAC(6′)-Ib-catalyzed acetylation of amikacin. **D.** Western blot showing total concentration of enzyme, in the wild-type background.

**Table S1.** – Strains used for this study.

| Strain | Relevant genotype | Source |
| --- | --- | --- |
| AB1157 | *thr-1*, *araC14*, *leuB6*(Am), DE(*gpt-proA*)62, *lacY1*, *tsx33*, *qsr<sup>-</sup>0*, *glnV44*(AS), *galK2*(Oc), LAM<sup>-</sup>, Rac-0, *hisG4*(Oc), *rfbC1*, *mgl-51*, *rpoS396*(Am), *rpsL31*(str<sup>R</sup>), *kdgK51*, *xylA5*, *mtl-1*, *argE3*(Oc), *thi-1* | (49) |
| BL21(DE3) | *fhuA2* [lon] *ompT gal* [dcm] Δ*hsdS* | (50) |
| TB25 | AB1157 derivative, *P<sub>lacq</sub>-lacI P<sub>Llac-s</sub>-dCas9 cat::ΔattB* | This work |
| FHcas1nc | TB25 derivative, *gRNA-Prom (20bp cod.)::argE* | This work |
| FHcas1 | TB25 derivative, *gRNA-Prom (20bp non-cod.)::argE* | This work |
| FHcas2nc | TB25 derivative, *gRNA-CDS (20bp cod.)::argE* | This work |
| LW3 | TB25 derivative, *gRNA-CDS (15bp cod.)::argE* | This work |
| TY1 | TB25 derivative, *gRNA-CDS (14bp cod.)::argE* | This work |
| LW2 | TB25 derivative, *gRNA-CDS (13bp cod.)::argE* | This work |
| LW1 | TB25 derivative, *gRNA-CDS (12bp cod.)::argE* | This work |
| TY2 | TB25 derivative, *gRNA-CDS (11bp cod.)::argE* | This work |
| TY3 | TB25 derivative, *gRNA-CDS (10bp cod.)::argE* | This work |
| FHcas2 | TB25 derivative, *gRNA-CDS (20bp non-cod.)::argE* | This work |
| FHcas3nc | TB25 derivative, *gRNA-Linker (18bp cod.)::argE* | This work |
| TY4 | TB25 derivative, *gRNA-Linker (14bp cod.)::argE* | This work |
| TY5 | TB25 derivative, *gRNA-Linker (11bp cod.)::argE* | This work |
| TY6 | TB25 derivative, *gRNA-Linker (10bp cod.)::argE* | This work |
| LW4 | TB25 derivative, *gRNA-Linker (9bp cod.)::argE* | This work |
| FHcas3 | TB25 derivative, *gRNA-Linker (15bp non-cod.)::argE* | This work |
| OD030 | *P<sub>Lac</sub>-mMaple kan::Δgalk* | This work |
| YHZ23 | *MATa his3Δ1 leu2Δ0 met15Δ0 LYS2 ura3Δ0 nup59-mNeonGreen-Nat* | This work |

**Table S2.**
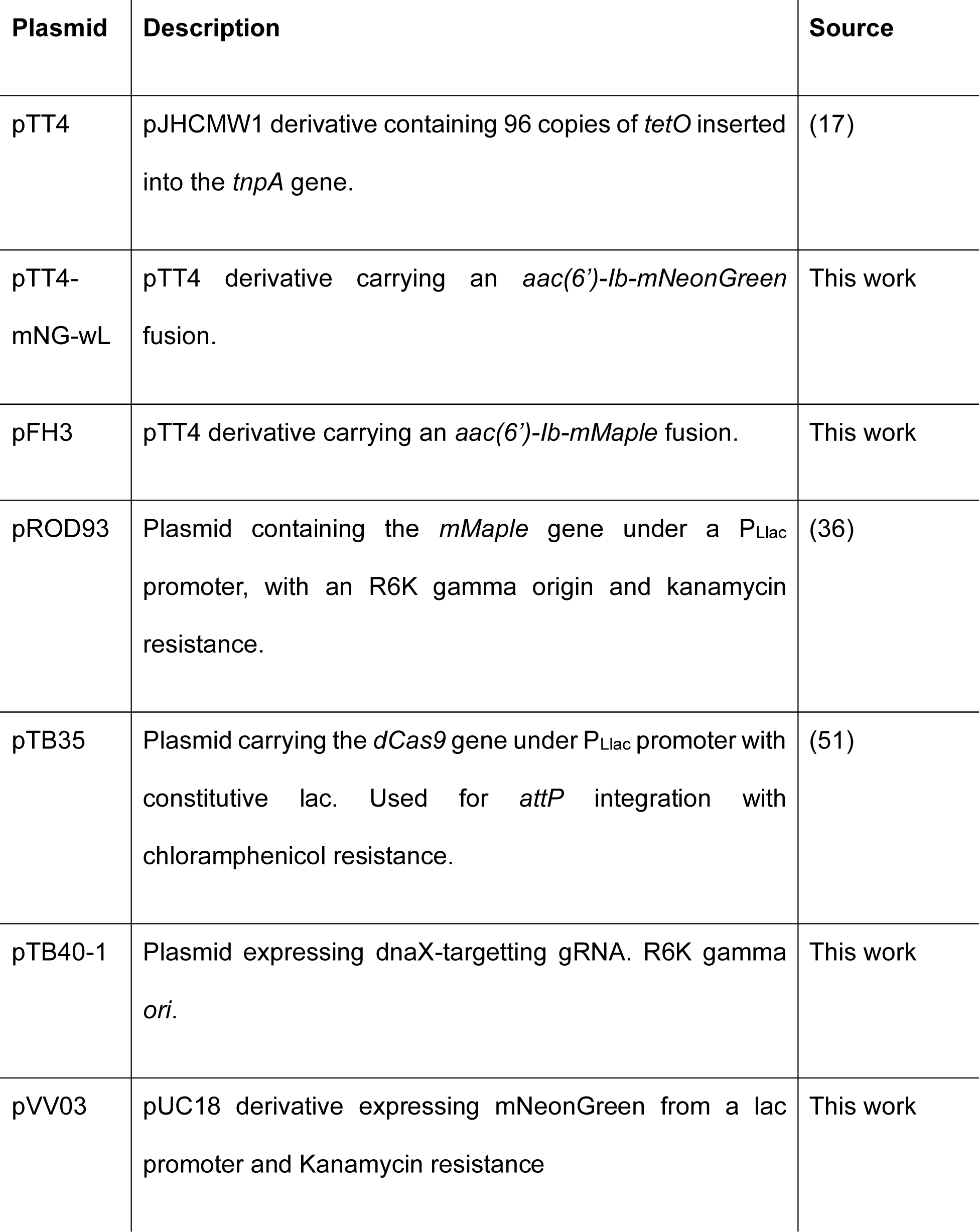
– Plasmids used in this study.

**Table S3.**
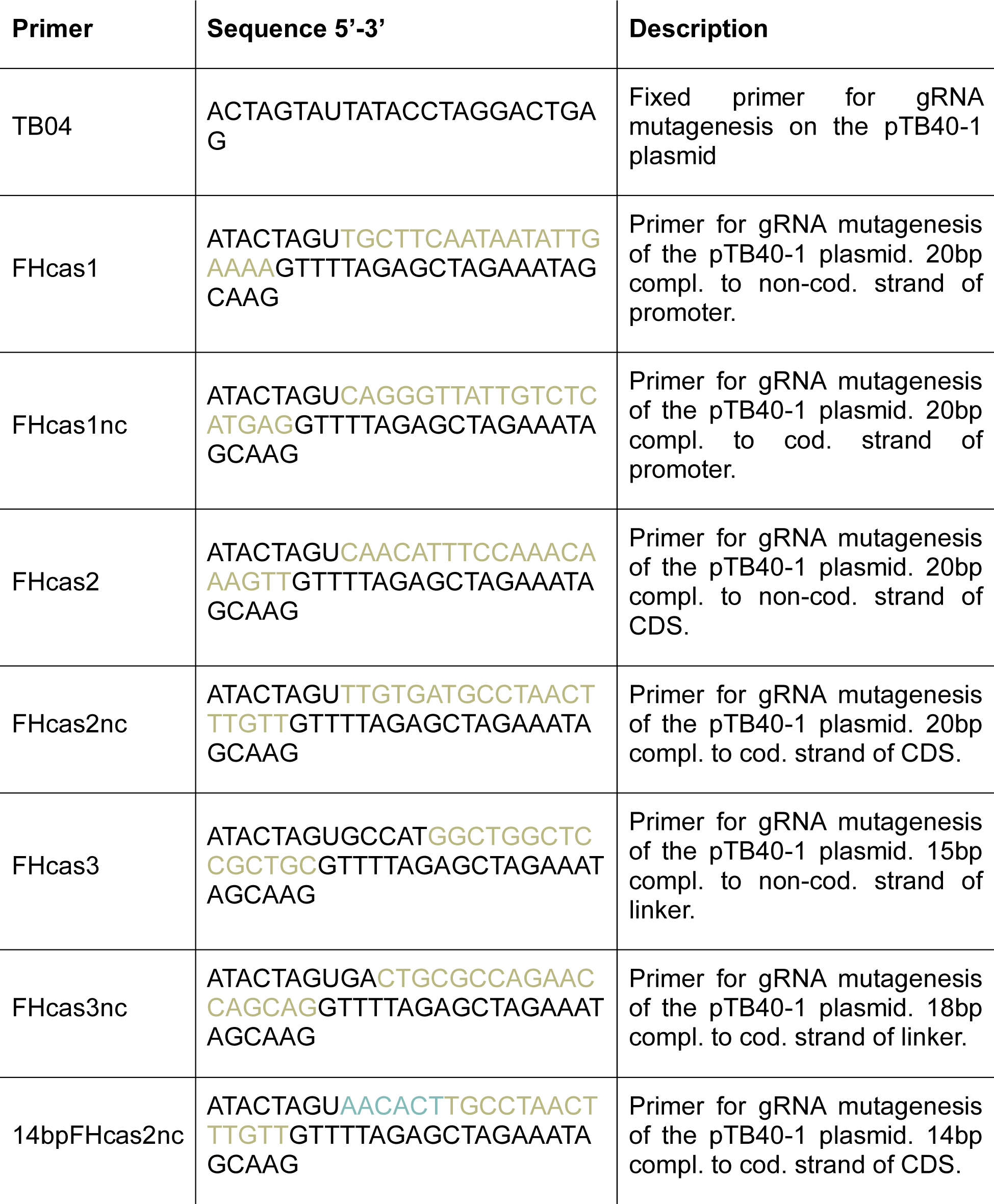
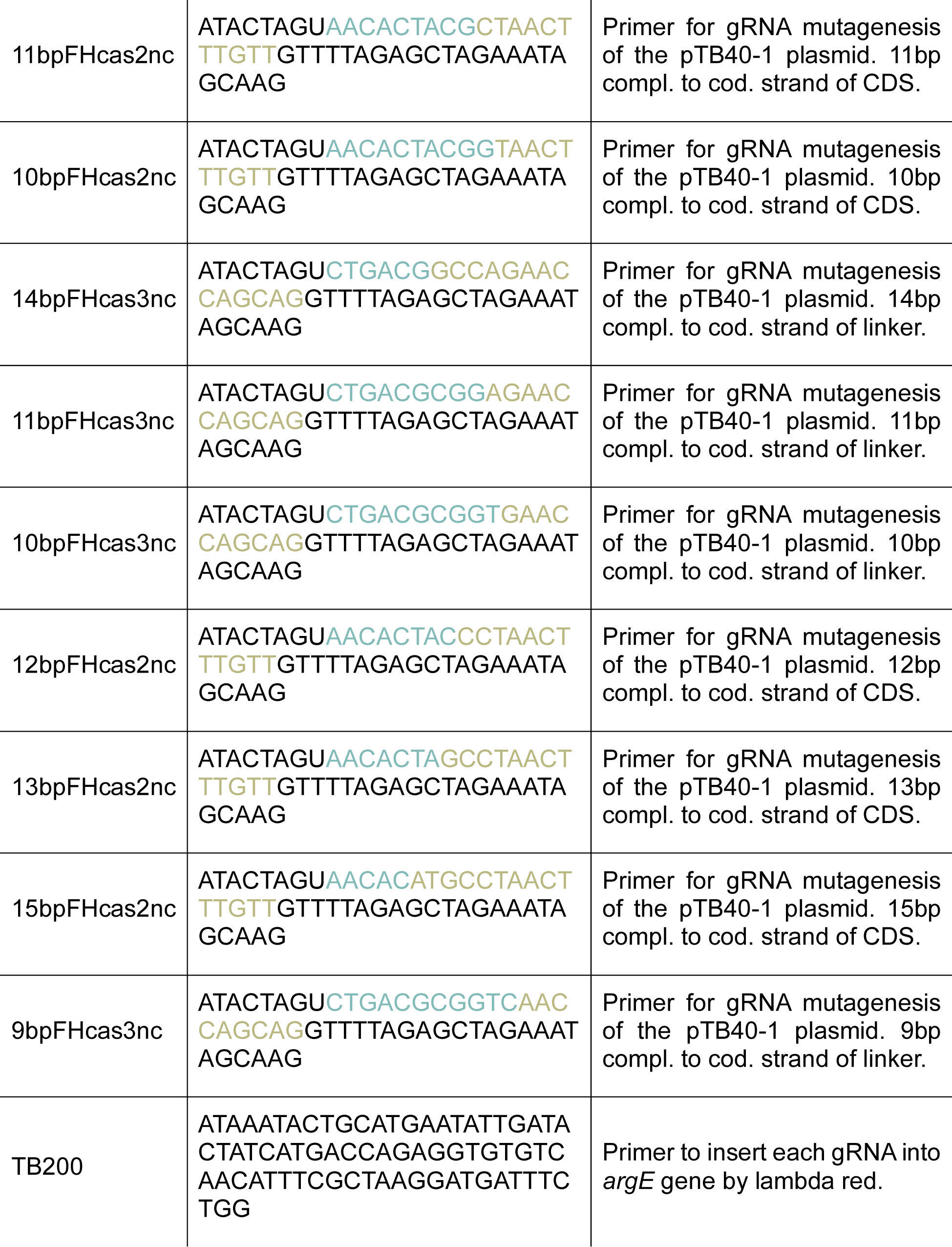
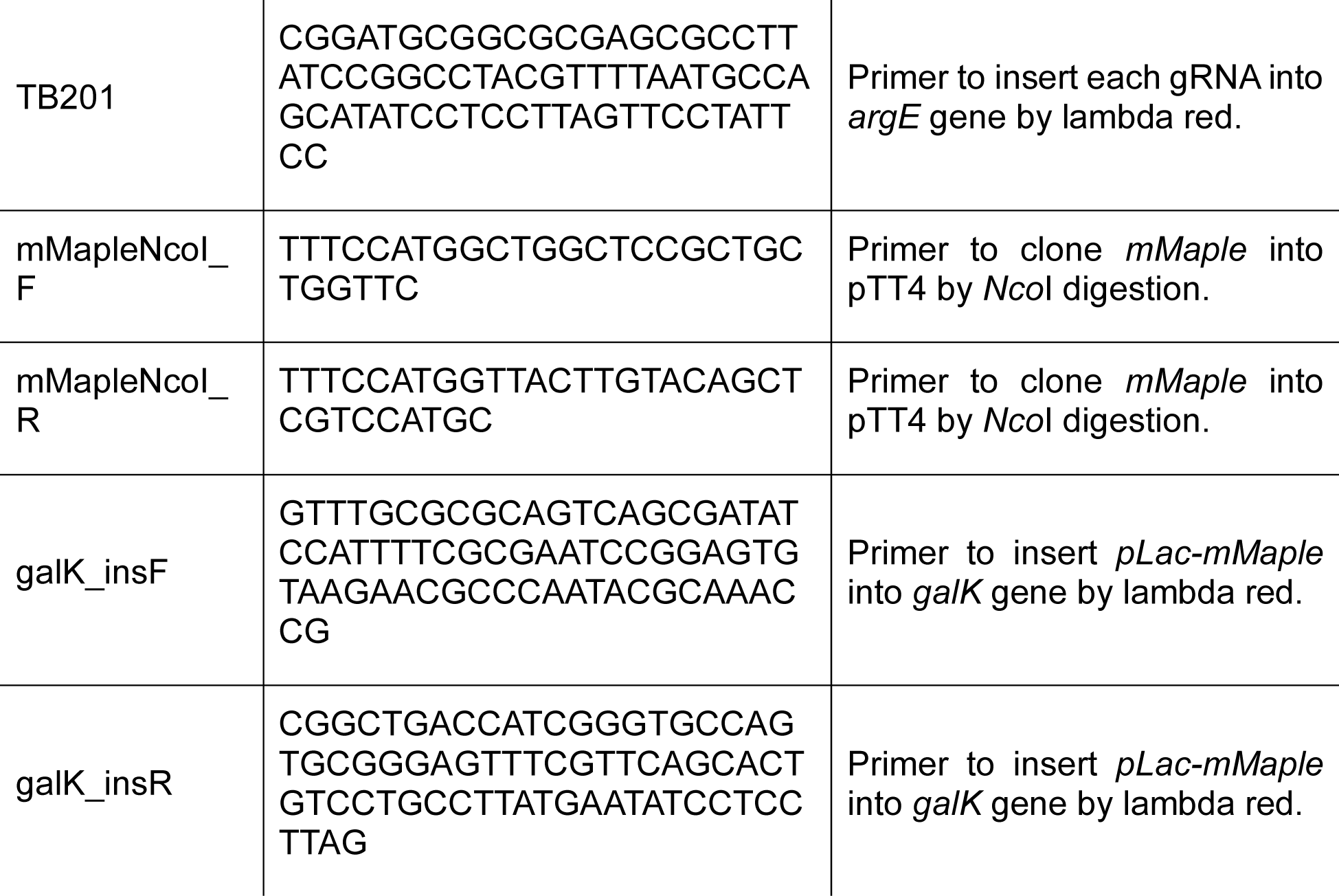
– Primers used in this work. The guide RNA sequences are highlighted in green, while the mutated base pairs are shown in blue. cod., coding, non-cod., non-coding, compl., complementary, CDS, coding sequence.

**Table S4.**
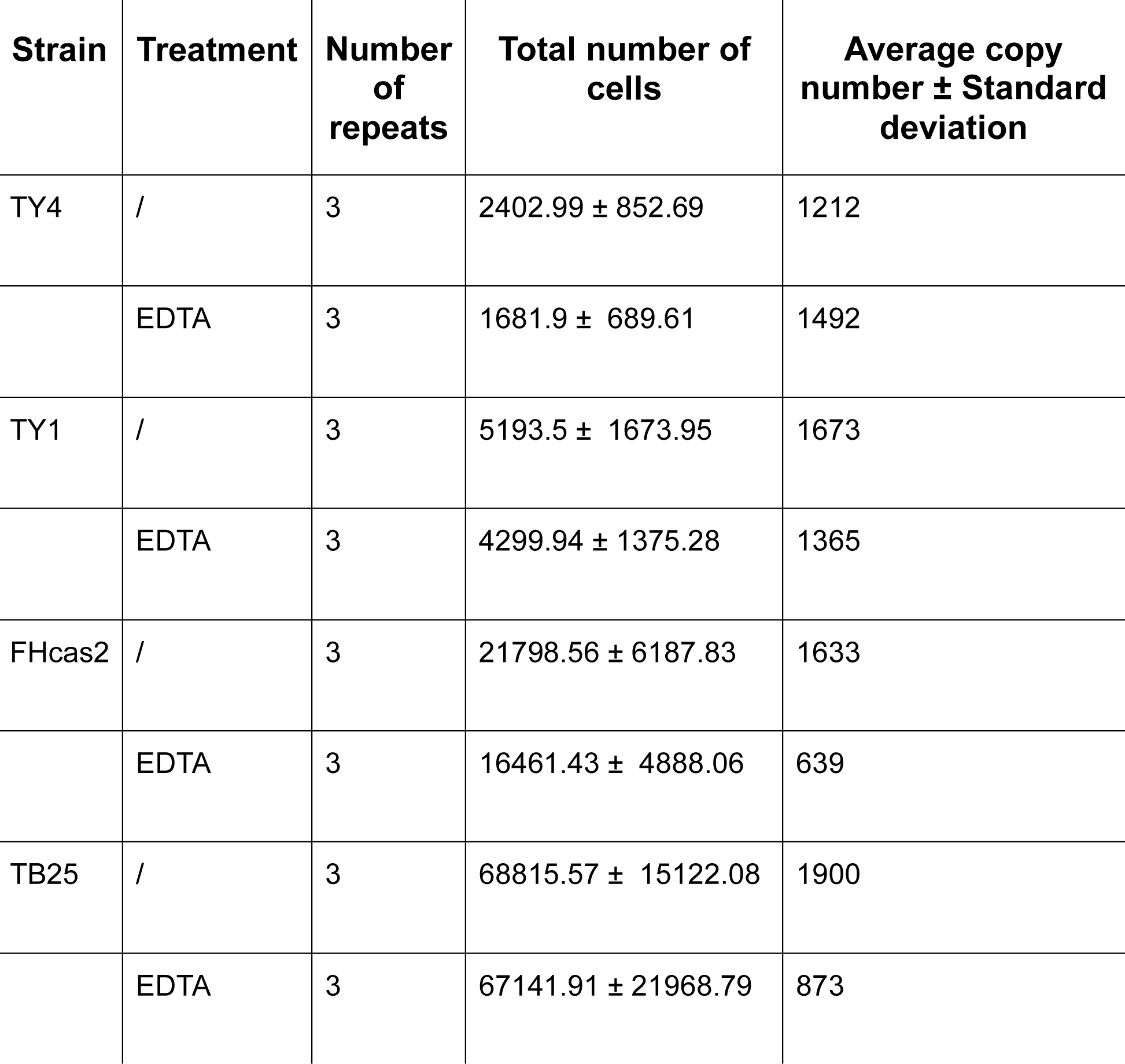
– Number of repeats and total cell count for Figure 3.D

| **Strain** | **Treatment** | **Number of repeats** | **Total number of cells** | **Average copy number ± Standard deviation** |
| --- | --- | --- | --- | --- |
| TY4 | / | 3 | 2402.99 ± 852.69 | 1212 |
| | EDTA | 3 | 1681.9 ± 689.61 | 1492 |
| TY1 | / | 3 | 5193.5 ± 1673.95 | 1673 |
| | EDTA | 3 | 4299.94 ± 1375.28 | 1365 |
| FHcas2 | / | 3 | 21798.56 ± 6187.83 | 1633 |
| | EDTA | 3 | 16461.43 ± 4888.06 | 639 |
| TB25 | / | 3 | 68815.57 ± 15122.08 | 1900 |
| | EDTA | 3 | 67141.91 ± 21968.79 | 873 |

